# Parabrachial-amygdala circuit cooperates with a posterior striatal area to drive opioid withdrawal aversion

**DOI:** 10.64898/2026.08.27.747629

**Authors:** Seung-Chan Lee, Kelsey Shimoda, Jesse Ross, Jensine Coudriet, Thomas Jhou, Satoshi Ikemoto

**Affiliations:** Neurocircuitry of motivation section, Behavioral Neuroscience Research Branch, Intramural Research Program, National Institute on Drug Abuse, National Institute of Health, Baltimore, Maryland 21224; Department of Neurobiology, University of Maryland School of Medicine, Baltimore, Maryland 21201

## Abstract

Opioid addiction treatment is often hampered by the severe dysphoria of opioid withdrawal, but withdrawal treatments are limited by incomplete understanding of brain mechanisms involved. One area frequently implicated in withdrawal symptoms is the central amygdala, whose capsular portion (CeC) is particularly strongly activated during withdrawal. Additionally, a ventral posterior striatal region that resides near CeC, the interstitial nucleus of the posterior limb of the anterior commissure (IPACc), is also activated as strikingly as CeC. However, it is still unknown how these regions are activated, nor whether their activation explains the high intensity of withdrawal dysphoria. Using RNAscope, we found that c-fos expression is induced in the parabrachial nucleus (PB), a key glutamatergic afferent of CeC, after precipitated morphine withdrawal. Chemogenetic inhibition of PB glutamatergic neurons (VG2^PB^) nearly eliminated withdrawal-induced CeC c-Fos, without affecting IPACc c-Fos, indicating these two nuclei are activated by distinct sources. Furthermore, VG2^PB^ inhibition markedly reduced somatic (jumping) and modestly reduced affective (place avoidance) withdrawal behavior. On the other hand, inhibition of CeC-projecting PB neuronal subtypes expressing calcitonin gene-related peptide (CGRP) or mu opioid receptor (MOR) reduced place avoidance without affecting jumping, indicating their specific role in withdrawal aversion. Strikingly, simultaneous inhibition of VG2^PB^ and posterior striatal region containing IPACc robustly reduced withdrawal-induced place avoidance much more than the modest effects of either inhibition alone, suggesting their cooperative action in driving aversion. Our data suggests that PB-CeC circuit and posterior striatal area constitute a cooperative system driving opioid withdrawal aversion.

## Introduction

Agonists at the mu opioid receptor (MOR) produce potent analgesic effects and euphoria, while discontinuation of these drugs in opioid-dependent individuals produces highly aversive withdrawal states [1]. The desire to avoid withdrawal distress drives individuals toward long term continuation of opioid use. Hence, mitigating the severity of opioid withdrawal would be an important step for the treatment of opioid use disorder. However, the neural basis of opioid withdrawal is still not well understood [2,3], in part because widespread MOR expression throughout the central nervous system [4] hinders identification of candidate neural circuits mediating opioid withdrawal. At the same time, this broad expression, especially in many areas related to motivation and emotion, suggests the possibility of recruitment of multiple aversive pathways related to different aspects of withdrawal aversiveness.

To identify neural substrates associated with withdrawal aversion, prior studies used c-Fos expression to identify activated neurons in brain areas after precipitated withdrawal [5]. One major region repeatedly identified is the central amygdala (CeA), particularly its capsular subregion (CeC) [5,6]. Given the CeA’s functional and anatomical association with the “extended amygdala (EA)”, a network implicated in stress responses [7,8], prior work also examined c-Fos within other EA regions. An area most strongly expressing c-Fos aside from the CeC was the caudal part of the interstitial nucleus of the posterior limb of the anterior commissure (IPACc), which borders the anterior aspect of the CeA [6]. Despite its particularly robust activation, this area’s role in opioid withdrawal has been largely neglected. Together with the CeC, it is poorly understood what circuits drive their activation, and how much such activations mediate physical and affective symptoms of opioid withdrawal.

Prior studies noted that CeC is critically involved in various pain processes [9–11], possibly in conjunction with its dense glutamatergic input from the parabrachial nucleus (PB) [12,13], another structure highly implicated in noxious stimuli processing [9,14–18]. This relationship suggests potential involvement of PB-CeC pain-related pathways during opioid withdrawal. In contrast, the IPACc, situated in the posterior ventral striatum, is located immediately anterior to the CeA, but exhibits strong connections to basal ganglia structures such as the globus pallidus (GP), substantia nigra reticulata (SNR) and midbrain dopaminergic neurons, while exhibiting sparser connections with other EA structures than the CeA [19–23]. The function of IPAC has only recently been linked to motivated behaviors and aversive processes [24–27], however, its contribution to the aversiveness of opioid withdrawal is completely unknown.

To understand CeC and IPACc roles in opioid withdrawal, we investigated circuit mechanisms of activation focusing on the impact of brainstem excitatory inputs, and whether these circuits contribute to the aversiveness of opioid withdrawal. Our data revealed that activation of the CeC, but not IPACc, is primarily driven by inputs from PB glutamatergic neurons. Furthermore, we found a reduction of withdrawal-induced conditioned place avoidance (CPA) after inhibition of PB glutamatergic neurons and subsets expressing calcitonin gene-related peptide (CGRP) or MOR. Strikingly, simultaneous inhibition of both PB glutamatergic neurons and a posterior striatal area that includes IPACc robustly reduced CPA more strongly than individual inhibitions of each region alone, suggesting a possibility that PB glutamatergic neurons and IPACc neurons cooperatively contribute to the aversiveness of withdrawal. This data identified a key role of PB neurons and their relationship with posterior striatal areas in driving aversiveness of opioid withdrawal.

## Materials and Methods

Detailed descriptions are provided in the Supplemental information.

### Animals and surgery

C57BL/6J, Vglut2-Cre and CGRP-Cre mice were purchased from Jackson Laboratory. MOR-Cre mice were provided by B. L. Kieffer. AAV injection surgery was performed as previously described [28]. All procedures were approved by the Animal Care and Use Committee of the National Institute on Drug Abuse.

### Naloxone-precipitated morphine withdrawal

Mice were injected subcutaneously twice a day with escalating doses of morphine sulfate pentahydrate (20, 20, 40, 40, 60, 60, 80, 80, 100, 100 mg/kg) over six days. Six hours after morphine injection on day 6, mice subcutaneously received naloxone hydrochloride dihydrate (0.19 mg/kg).

### Detection of c-Fos expression

c-Fos immunohistochemistry was performed as previously described with modifications [29].

### RNAscope in situ hybridization

Brains were removed with fresh frozen method. RNAscope fluorescence multiplex v1 and v2 kits (Advanced Cell Diagnostics) were used.

#### Locomotor activity assay

The number of IR beam breaks during 15 min in mouse chambers was counted as readout of locomotor activity.

### Withdrawal-induced conditioned place avoidance

CPA box contains left, right and middle compartments. The left and right compartments are distinguished by visual, tactile and olfactory cues: mice were habituated to the CPA box for about 5 min on day -1. On day 0, each mouse was allowed to explore the box freely for 15 min. On day 5 of morphine injection schedule, six hours after morphine injection, mice were confined in the left chamber after receiving saline for 20 min. On day 6, six hours after morphine injection, mice were confined in the right side after receiving naloxone for 20 min. Jumping incidents were counted manually offline. On day 7, mice were allowed to explore the entire box freely for 15 min. The level of CPA was accessed by CPA score (= Time_pre_-Time_post_) and normalized CPA score. After the CPA test on day 7, with some mice, additional morphine injections (60, 80, 100, 100 mg/kg) over 3 consecutive days were made before withdrawal induction and brain extraction for c-Fos assay.

### Statistical Analysis

Default statistical test is the Student’s t test (means ± SEM), when data passed normality test. Otherwise, non-parametric tests (Mann-Whiteney for unpaired test, Wilcoxon signed-rank test for paired test) were used (median ± IQR). All tests were two sided.

## Results

### Naloxone-precipitated morphine withdrawal increases c-Fos in the CeC and IPACc

We used a naloxone-precipitated morphine withdrawal procedure to investigate the roles of the CeA and neighboring regions in multiple aspects of opioid withdrawal. Morphine was administered subcutaneously twice daily over six days with escalating doses [30] (Fig. 1A). Six hours after the last morphine administration on day 6, the opioid receptor antagonist naloxone was injected to precipitate withdrawal symptoms. This procedure robustly increased c-Fos (a marker of neuronal activation) protein expression in the CeA (Fig. 1B-C). Specifically, c-Fos positive (c-Fos+) neurons were dense in the CeC, while the rest of CeA had low to moderate expression levels. We also observed robust c-Fos expression in the IPACc adjacent to the anterior part of CeA (Fig. 1C), consistent with a previous report in rats [6]. By contrast, naloxone injections in chronic saline-injected mice did not induce such c-Fos expressions (Fig. 1B-C). Hence, opioid withdrawal robustly activates CeC and IPACc neurons.

**Figure 1.**
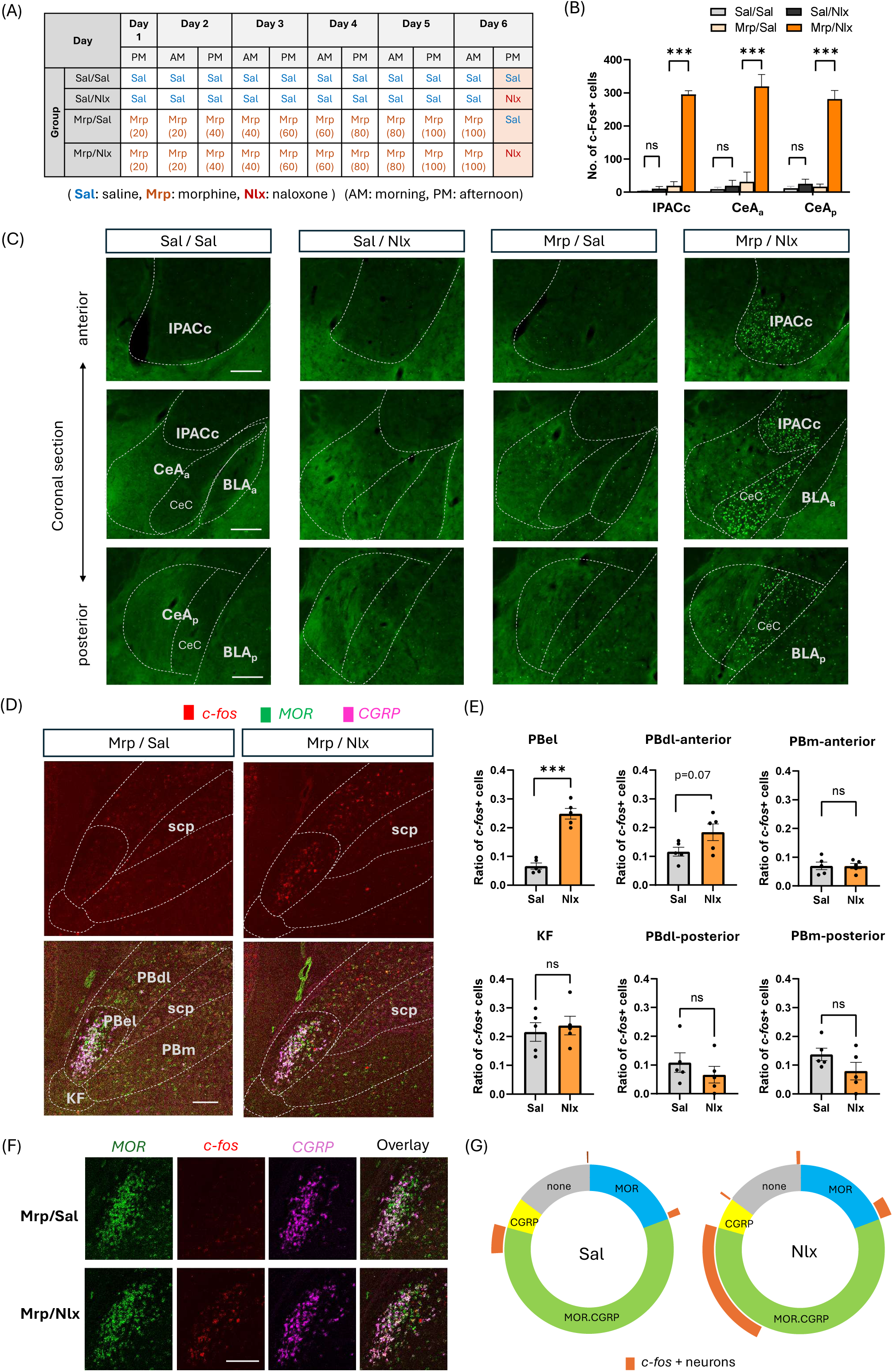
Activation of CeA, IPACc and PB during opioid withdrawal. **(A)** Timeline of c-Fos experiment with precipitated morphine withdrawal. Four groups received either morphine (Mrp) or saline (Sal) injections (s.c) twice daily over 6 days followed by naloxone (Nlx) or Sal injection 6 hours after the last injection of Mrp or Sal on day 6. Numbers in parentheses are morphine sulfate pentahydrate doses (mg/kg). **(B)** Numbers of c-Fos+ neurons per 50 µm section in IPACc, anterior CeA (CeA_a_), and posterior CeA (CeA_p_) regions after morphine withdrawal. *** p < 0.001 (Tukey tests preceded by significant group differences by one-way ANOVAs: F(3,8)=720, p < 0.001 for IPACc; F(3,8)=110, p < 0.001 for CeA_a_; F(3,8)=212, p < 0.001 for CeA_p_). **(C)** Representative images of c-Fos immunoreactivity in coronal brain sections at 3 anterior-posterior (AP) levels from the 4 experimental groups in (A). Anterior, mid and posterior regions correspond approximately to AP -0.4, -0.8, and -1.2mm from bregma in Franklin and Paxinos’s mouse brain atlas [56], respectively. **(D)** Representative mRNA signals of *c-fos*, MOR and CGRP in PB area (coronal section) after morphine withdrawal treatment in two conditions: Mrp/Sal (left) and Mrp/Nlx (right). Top: *c-fos* signals only. Bottom: overlaid images of *c-fos*, MOR and CGRP gene signals. **(E)** Ratio of *c-fos*+ neurons (*c-fos*+ cells / total DAPI+ cells) in six PB subregions of Mrp/Sal (gray bar) and Mrp/Nlx (orange bar) groups. *** p<0.001 (two-tailed T-test). **(F)** Representative mRNA signals of *c-fos*, MOR and CGRP in PBel region. **(G)** Distribution of *c-fos*+ neurons among PBel neuron types classified by MOR and CGRP expression in Mrp/Sal (left) and Mrp/Nlx (right) groups. Abbreviations: CeA_a_, anterior CeA; CeA_p_, posterior CeA; CeC, capsular region of CeA; BLA_a_, anterior basolateral amygdala; BLA_p_, posterior basolateral amygdala, PBel, external lateral region of PB; PBdl, dorsolateral region of PB; PBm, medial region of PB; KF, Kolliker-Fuse region; scp, superior cerebellar peduncle. Scale bars, 200 µm.

### Naloxone-precipitated withdrawal increases *c-fos* mRNA in the PB

To investigate the neuronal pathways that activate CeC and IPACc neurons during withdrawal, we examined the PB as an important candidate upstream structure that could contribute to the neuronal activation. The PB sends glutamatergic projections to the CeA and nearby EA structures, and is one of the regions that express the highest level of MOR in the brain [4]. In addition, MOR+ PB neurons project to the CeA [31], with the CeC receiving particularly strong projections from the external lateral part of the PB (PBel) [9,12] that are implicated in negative affect of pain [9,14,32].

We first examined whether precipitated withdrawal significantly activates PB neurons. Instead of c-Fos protein, we measured *c-fos* mRNA, due to our findings that morphine itself may induce c-Fos protein expression that is observable in the PB 7.5 hours after morphine injections, without naloxone treatment (Fig. S1). Hence, we examined *c-fos* mRNA, which has a markedly shorter half-life than c-Fos protein [33,34] to minimize compounding signals produced before withdrawal induction. We observed low levels of *c-fos* mRNA in morphine-treated mice without naloxone precipitation, while naloxone injections significantly increased *c-fos* mRNA expression in the PBel, but not other PB sub-regions (Fig. 1D-E). However, we cannot rule out subthreshold effects in these other sub-regions, such as the dorsolateral region of the PB (PBdl), which displayed an insignificant trend toward increased *c-fos* mRNA.

Because CGRP+ neurons in the PBel provide key inputs to the CeA [9,12], we then examined whether MOR gene (*Oprm1*) expression overlaps with CGRP gene (*Calca*) expression in the PBel and whether these populations are activated by naloxone-precipitated withdrawal. Using RNAscope, we found MOR+ neurons throughout the PB, while the majority of CGRP+ neurons are confined in the PBel (Fig. 1D). Overall, 76% of MOR+ neurons in the PBel were CGRP+, while 91% of CGRP+ neurons were MOR+ in this region (Fig. 1F-G). Naloxone-precipitated withdrawal induced *c-fos* in 28% of PBel neurons, while conversely 97% of *c-fos*+ PBel neuron were MOR+, and 82% of *c-fos*+ PBel neurons were CGRP+. In addition, 81% of PBel *c-fos*+ neurons were double positive for both MOR and CGRP (Fig. 1F-G). These data support the idea that opioid withdrawal activates PBel neurons, particularly those co-express MOR and CGRP, which then activate CeA and nearby neurons.

### Inhibitions of MOR- and CGRP-expressing PB neurons decrease CeC c-Fos and place avoidance induced by precipitated withdrawal

The abovementioned results led us to examine whether specific subtypes of PB neurons activate CeC and IPACc neurons and induce negative affect triggered by opioid withdrawal. We used a DREADD procedure to inhibit PB neurons and Cre mouse lines to address the cell types. Male mice were used throughout the study, because we found that withdrawal-induced conditioned place avoidance (CPA) that was consistently measurable in both males and females, but was stronger and less variable in males (Fig. S2).

We first tested effects of inhibiting PB MOR+ neurons (MOR^PB^) on gross locomotor activity using an AAV-hsyn-DIO-hM4Di injected into the PB of MOR-Cre mice (Fig. 2A-C). Inhibition of MOR^PB^ did not affect baseline locomotor activity (Fig. 2D). Four days after the locomotion test, mice began to receive escalating doses of morphine (2x a day over 6 days). During the final two days of morphine injections, mice received a place conditioning procedure (Fig. 2A) in which one side of the CPA apparatus was paired with saline, while the other side was paired with naloxone injection. During withdrawal, we observed intense jumping, a characteristic physical symptom of opioid withdrawal in mice, and the withdrawal-induced jumping did not decrease with the inhibition of MOR^PB^ by CNO (Fig. 2E). On the next day, CPA was measured without any injection of CNO or saline. We first calculated the CPA score, which is the difference in times that mice spent in naloxone-paired side between before and after the conditioning. We also calculated normalized CPA score by dividing the raw CPA score by the pre-conditioning time in the naloxone-paired chamber to control for variability of the pre-conditioning time values [35]. MOR^PB^ inhibition with CNO during withdrawal decreased both raw and normalized CPA scores (Fig. 2F-G). CNO treatment in the absence of DREADD did not affect withdrawal-induced CPA in MOR-Cre mice (Fig. S3A). The data suggest that MOR^PB^ neurons play a role in negative affect induced by opioid withdrawal, but do not contribute to jumping.

**Figure 2.**
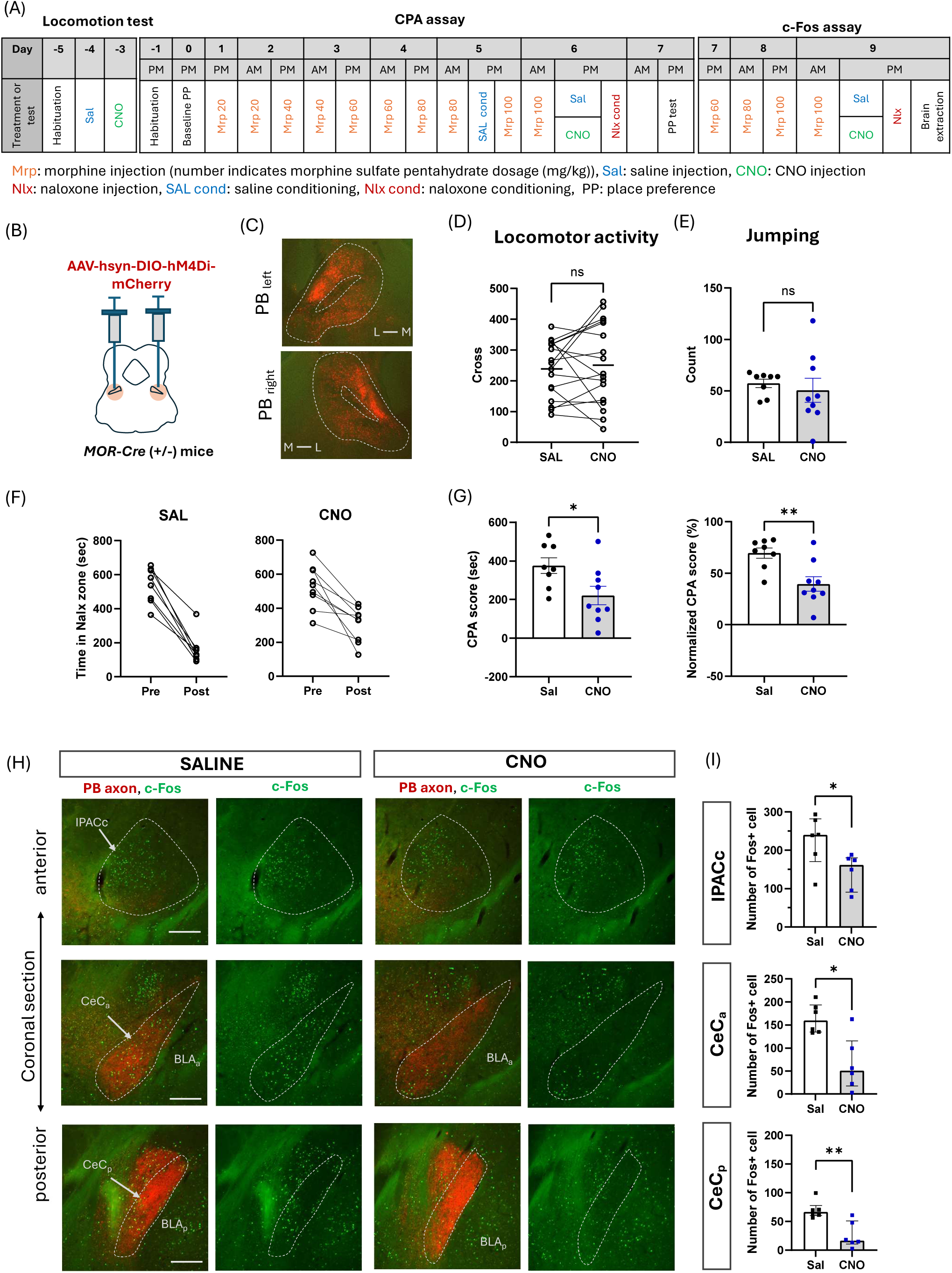
MOR^PB^ inhibition decreased withdrawal-induced place avoidance and c-Fos counts in CeC and IPACc. **(A)** Experimental timeline for locomotor, precipitated withdrawal-induced CPA and c-Fos tests. **(B)** MOR-Cre mice received bilateral injections of an AAV-hsyn-DIO-hM4Di-mCherry into the PB. **(C)** Coronal sections showing representative expressions of hM4Di-mCherry in MOR^PB^. **(D, E)** Chemogenetic inhibition of MOR^PB^ did not affect baseline locomotor activity. (paired t-test) and withdrawal-induced jumping (Mann-Whitney test). **(F)** Time spent in naloxone-paired compartment before and after conditioning. **(G)** MOR^PB^ inhibition significantly decreased both CPA score (left) and normalized CPA score (right). (two-tailed t-test). **(H)** Coronal sections showing representative c-Fos expression in IPACc (top), CeC_a_ (anterior CeC, middle), and CeC_p_ (posterior CeC, bottom) of saline- and CNO-treated mice. Dotted lines outline IPACc, CeC_a_, and CeC_p_. Green dots indicate c-Fos immunoreactivity, while red signals indicate mCherry-labeled axons of MOR^PB^. See Figure 1 legend for abbreviations. **(I)** Number of c-Fos+ neurons per section in IPACc, CeC_a_, CeC_p_ of saline- and CNO-treated mice. (Mann-Whitney test). * p<0.05, ** p<0.01 Scale bar, 200 µm.

To understand how MOR^PB^ inhibition affects withdrawal-induced activations of the CeC and IPACc, we examined c-Fos expression after inducing withdrawal with additional morphine administrations in a subset of these animals (Fig. 2A). This second withdrawal procedure produced a similar pattern of c-Fos expression in CeC and IPACc (Fig 2H-I) as those of the withdrawal (in mice lacking DREADD expression) described in Figure 1, albeit with fewer numbers of c-Fos+ neurons. DREADD-mediated inhibition of MOR^PB^ significantly decreased c-Fos expression in CeC compared to saline controls (Fig. 2H-I). Interestingly withdrawal-induced c-Fos in IPACc was also mildly reduced by CNO treatment, even though MOR^PB^ send no or very weak projections to the IPACc area (Fig. 2H-I). Together, these results suggest that MOR^PB^ activate CeC neurons possibly via monosynaptic pathways while also contributing to IPACc activation indirectly via polysynaptic pathways during opioid withdrawal.

We next examined the roles of PB CGRP+ neurons (CGRP^PB^) using CGRP-Cre mice. We found that bilateral inhibition of CGRP^PB^ did not affect baseline locomotor activity or withdrawal-induced jumping (Fig. 3A-D). Although CGRP^PB^ inhibition did not significantly reduce raw CPA scores, it reduced normalized CPA scores (Fig. 3E and S4), suggesting a modest effect. Again, CNO had no effect on withdrawal-induced CPA without DREADD expression (Fig. S3B). In addition, CGRP^PB^ inhibition reduced c-Fos in CeC, but less strongly compared to MOR^PB^ inhibition, while IPACc c-Fos was not affected (Figs. 3F and S4).

**Figure 3.**
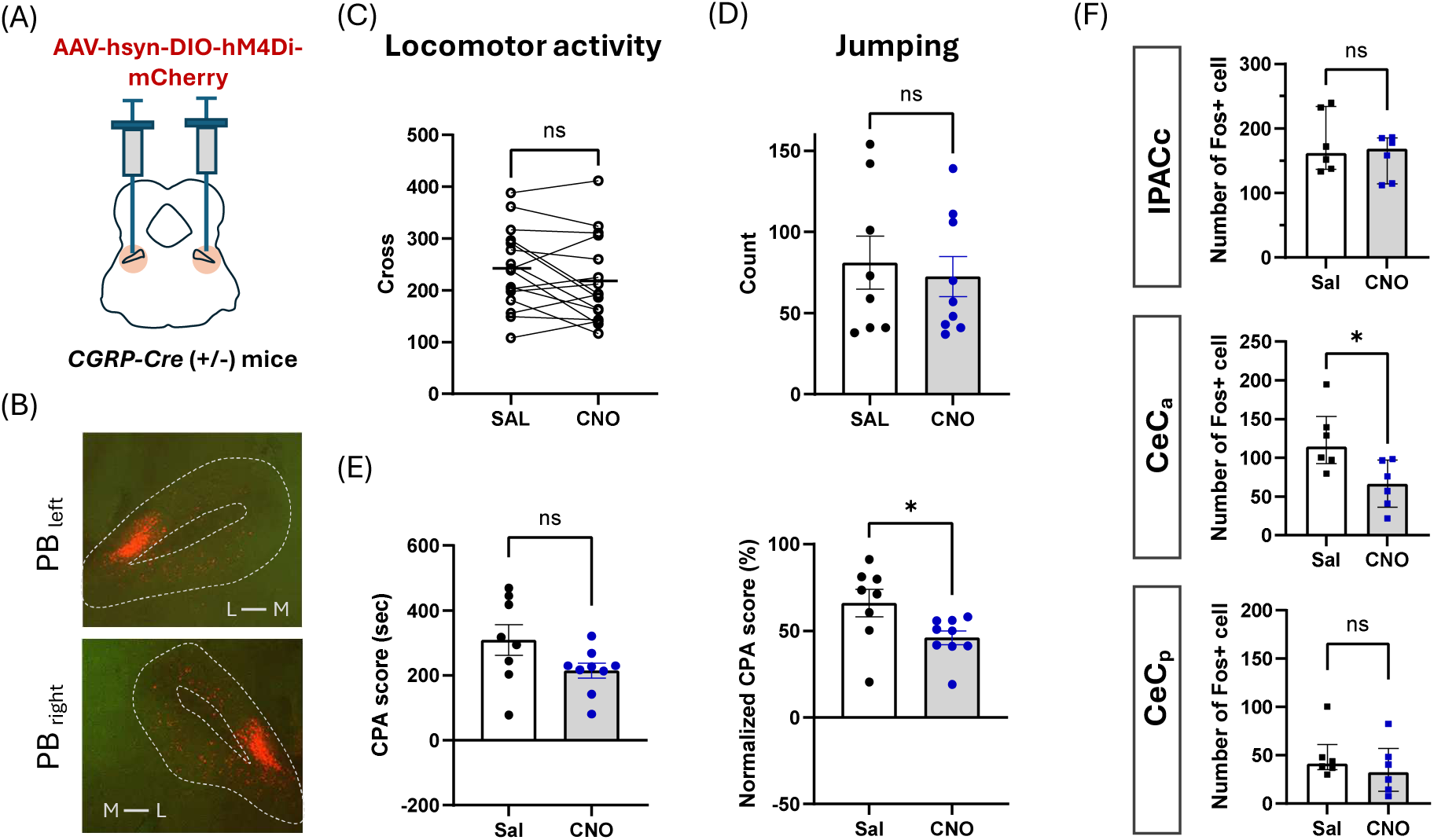
CGRP^PB^ inhibition decreases withdrawal-induced place avoidance and c-Fos counts in CeC. **(A)** CGRP-Cre mice received bilateral injections of an AAV-hsyn-DIO-hM4Di-mCherry into the PB. **(B)** Coronal sections of the PB showing representative expressions of hM4Di-mCherry in CGRP^PB^. **(C, D)** Chemogenetic inhibition of CGRP^PB^ did not affect baseline locomotor activity (paired t-test), or withdrawal-induced jumping (two-tailed t-test). **(E)** CGRP^PB^ inhibition did not significantly decrease CPA score (left), but decreased normalized CPA score (right) (two-tailed t-test). **(F)** Number of c-Fos+ neurons per section in IPACc, CeC_a_, CeC_p_ of saline- and CNO-treated mice. (Mann-Whitney). * p<0.05. Scale bar, 200 µm.

These data together suggest that precipitated withdrawal activates MOR^PB^ and CGRP^PB^, which, in turn, activate EA and contribute to negative affect, but without influencing physical symptoms such as jumping. Because a substantial portion of PB neurons still do not contain MOR or CGRP, we next inhibited PB neurons in a less-selective manner.

### Inhibition of PB Vglut2-expressing neurons decreases CeC c-Fos and place avoidance induced by precipitated withdrawal

We next examined the role of PB glutamatergic neurons, the major neuron type of the PB [36], in CeC and IPACc activation and withdrawal-induced negative affect. AAV-hsyn-DIO-hM4Di was injected into the PB in Vglut2-Cre mice, to selectively target PB glutamatergic neurons (Fig. 4A-B). Interestingly, we observed notable differences in addition to similarities with MOR^PB^ or CGRP^PB^ inhibitions. Inhibition of VG2^PB^ neurons did not significantly affect locomotor activity (Fig. 4C), although we observed that mice walked in an awkward manner after VG2^PB^ inhibition (observation not quantified). VG2^PB^ inhibition, unlike MOR^PB^ or CGRP^PB^ inhibitions, robustly decreased withdrawal-induced jumping (Fig. 4D), and reduced the normalized CPA score without significantly affecting raw CPA scores (Fig. 4E-F). Again, CNO itself did not affect withdrawal-induced CPA in Vglut2-Cre mice lacking DREADD expression (Fig. S3C). Furthermore, VG2^PB^ inhibition strongly suppressed withdrawal-induced c-Fos expression in the CeC (Fig. 4G-H), while not affecting IPACc c-Fos. This suggests that withdrawal-induced activation of VG2^PB^ is critical for cellular activation of CeC, but does not affect IPACc activation.

**Figure 4.**
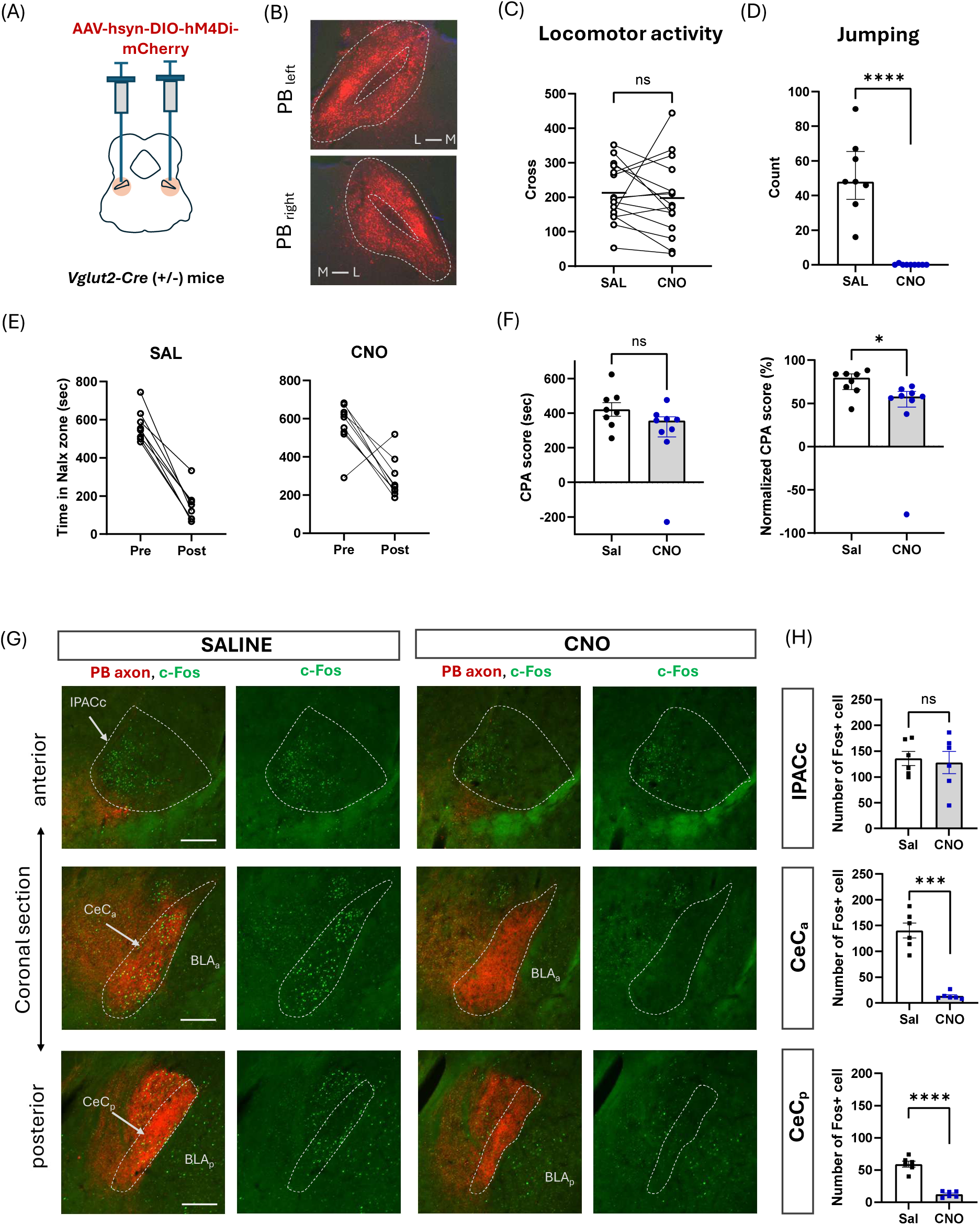
VG2^PB^ inhibition decreases withdrawal-induced jumping, place avoidance and c-Fos counts in CeC. **(A)** Vglut2-Cre mice received bilateral injections of an AAV-hsyn-DIO-hM4Di-mCherry into the PB. **(B)** Coronal sections showing representative expressions of hM4Di-mCherry in VG2^PB^. **(C, D)** Chemogenetic inhibition of VG2^PB^ did not affect baseline locomotor activity (paired t-test), but decreased withdrawal-induced jumping (Mann-Whitney test). **(E)** Time spent in naloxone-paired compartment before and after conditioning. **(F)** VG2^PB^ inhibition did not significantly decrease CPA score (left), but decreased normalized CPA score (right) (Mann-Whitney test). **(G)** Coronal sections showing representative c-Fos expressions of saline- and CNO-treated mice. See Figure 1C and 2H legend for additional description. **(H)** Number of c-Fos expressing neurons per section in IPACc, CeC_a_, CeC_p_ of saline- and CNO-treated mice (two-tailed t-test). * p < 0.05, *** p<0.001, **** p<0.0001. Scale bar, 200 µm.

Notably, VG2^PB^ inhibition appears less effective than MOR^PB^ inhibition in reducing both CPA and IPACc c-Fos expression, even though Vglut2 is expressed in the vast majority of PB neurons, including MOR^PB^. These findings may reflect the functionally and molecularly heterogeneous nature of the VG2^PB^ population [36–40], and raise the possibility that simultaneous inhibition of all subtypes of VG2^PB^ could mask the effects of inhibiting specific subtypes of VG2^PB^ such as MOR^PB^.

### Dual inhibition of PB Vglut2-expressing neurons and IPACc area neurons strongly decreases withdrawal-induced place avoidance

Our PB inhibition experiments demonstrated a critical influence of PB neuronal activity on CeC activation. In particular, CeC c-Fos expression was nearly eliminated by VG2^PB^ inhibition, while CPA was only modestly reduced. Hence, blockade of CeC activation may be insufficient to block the majority of withdrawal-induced aversion as measured by place avoidance, but it is not known what other structures may contribute additionally. Because IPACc activation was only minimally affected by our PB inhibitions, we next asked whether IPACc neurons could play such a role.

In order to inhibit IPACc neurons, we injected AAV-CaMKIIα-hM4Di into the IPACc area, taking advantage of high CaMKIIα expression in striatum [41,42], including IPACc [43] (Fig. 5A-B). This approach combined with CNO treatment silenced IPACc c-Fos expression induced by precipitated withdrawal (Fig. 5B and S5), although hM4Di expression also spread beyond the borders of the relatively small IPACc and into nearby regions, such as the anterior portion of CEA and striatal regions dorsal to IPACc. This inhibition of IPACc and adjacent areas did not affect baseline locomotor activity (Fig. 5C), but slightly increased withdrawal-induced jumping (Fig. 5D). Also, IPACc area inhibition moderately reduced the normalized but not raw CPA score (Fig. 5E-F and S3D-E). Notably, although hM4Di expression did spread to CEA, this was mostly restricted to its anterior portions, and its extent did not correlate with either raw or normalized CPA scores (Fig. S6).

**Figure 5.**
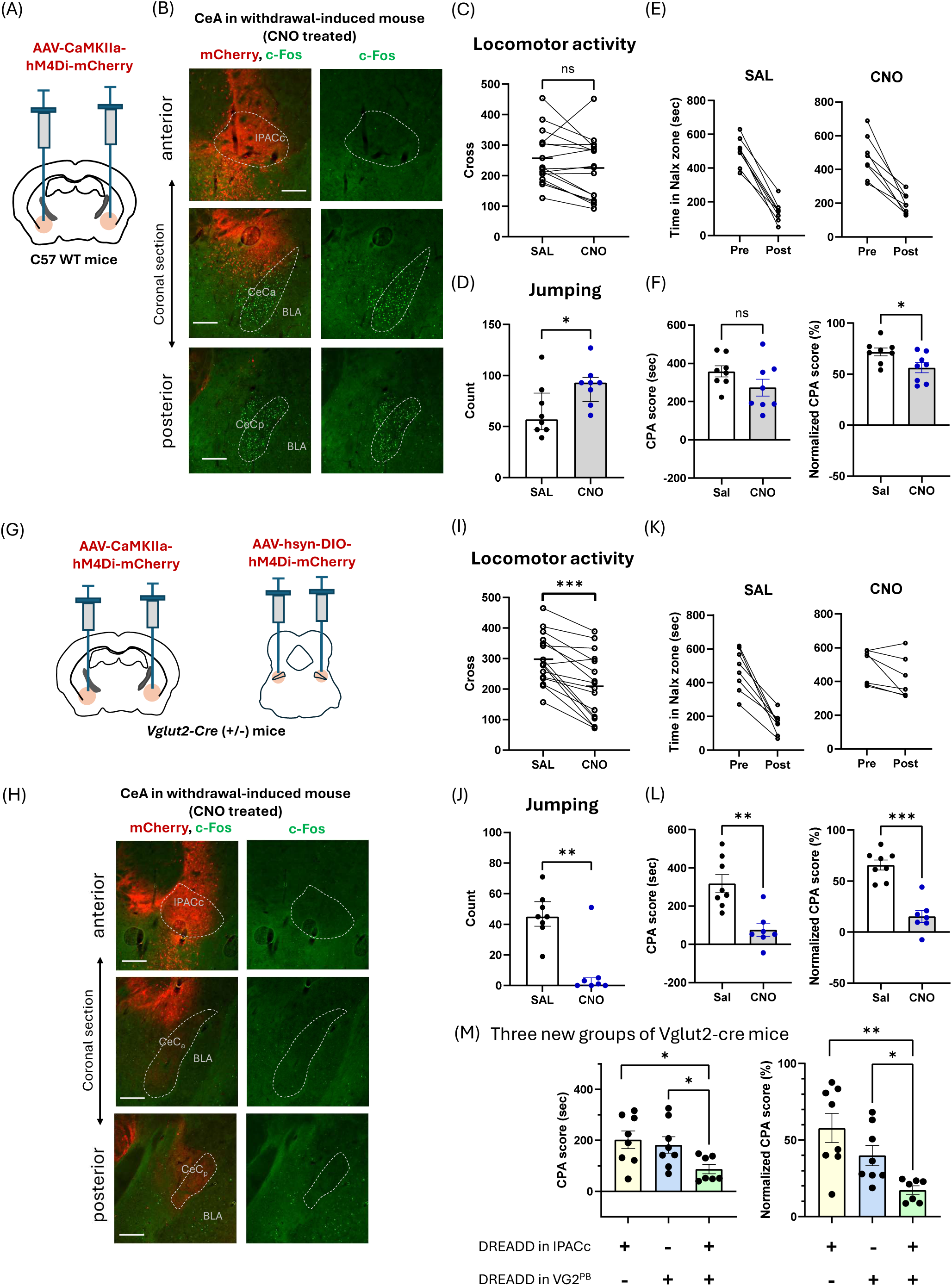
Dual inhibition of VG2^PB^ and IPACc area neurons produces strong reduction of withdrawal-induced place avoidance. **(A)** WT mice received bilateral injections of an AAV-CaMKIIa-hM4Di-mCherry into the IPACc area. **(B)** Coronal sections showing representative withdrawal-induced c-Fos expressions in CeA area of CNO-treated mice. See Figure 1 legend for abbreviations. **(C, D)** Chemogenetic inhibition of IPACc area neurons did not affect locomotor activity (paired t-test), and increased withdrawal-induced jumping. * p < 0.05 (Mann-Whitney test). **(E)** Time spent in naloxone-paired compartment before and after conditioning. **(F)** Inhibition of IPACc area neurons did not significantly decrease CPA score (left), but decreased normalized CPA score (right) (two-tailed t-test). **(G)** Schematic of bilateral injections of the AAVs for simultaneous dual inhibition of IPACc area neurons and VG2^PB^ in Vglut2-Cre mice. **(H)** Coronal sections showing representative withdrawal-induced c-Fos expressions in CeA area of CNO-treated mice. See Figure 2 legend for abbreviations. **(I, J)** Dual inhibition of IPACc area neurons and VG2^PB^ mildly decreased locomotor activity (paired t-test), and decreased withdrawal-induced jumping (Mann-Whitney test). **(K)** Time spent in naloxone-paired compartment before and after conditioning. **(L)** Dual inhibition of IPACc area neurons and VG2^PB^ significantly decreased both CPA score (left) and normalized CPA score (right) (two-tailed t-tests). **(M)** Three additional groups of Vglut2-cre mice received bilateral injections of inhibitory DREADD AAVs either in PB only, IPACc area only, or both, followed by a test of CPA. All 3 groups received CNO injection on day 6, 30 min prior to withdrawal induction to inhibit DREADD-expressing neurons. Withdrawal-induced CPA scores were lower after simultaneous inhibition of both IPACc area and VG2^PB^ relative to inhibitions of each region alone; CPA score (left), normalized CPA score (right) (two-tailed t-test with Holm Bonferroni multiple comparison correction). * p < 0.05, ** p < 0.01, *** p<0.001. Scale bar, 200 µm.

Finally, we tested the effects of simultaneously inhibiting IPACc area neurons and VG2^PB^ neurons, by injecting AAV-CaMKIIα-hM4Di into the IPACc area and AAV-hsyn-DIO-hM4Di into the PB of Vglut2-Cre mice. In these mice, CNO treatment effectively silenced withdrawal-induced c-Fos expressions in both IPACc and CeC (Fig. 5G-H and S5), while significantly decreasing baseline locomotion (Figure 5I) and withdrawal-induced jumping (Fig. 5J). Importantly, the dual inhibitory manipulation strongly decreased CPA (Fig. 5K-L) unlike the inhibition of VG2^PB^ or IPACc area neurons separately observed above. However, in these experiments, we had not explicitly planned a comparison between the dual-inhibition group and the two single-inhibition groups. Hence, we next tested three new groups of Vglut2-Cre mice (Fig. 5M) receiving DREADD inhibition of either VG2^PB^ alone, IPACc area neurons alone, or both regions combined, and compared the CPA scores between them. As hypothesized, this experiment showed that inhibiting two regions produced a stronger reduction in withdrawal-induced CPA than inhibition of either region alone. We again saw minor spread into the CEA, but the degree of spread was uncorrelated with CPA scores (Fig. S6). These data together suggest the possibility that IPACc area neurons participate in mediating aversive effect of opioid withdrawal and can intensify aversiveness in concert with VG2^PB^ activity.

## Discussion

We found that parabrachial glutamatergic neurons and their subpopulations expressing MOR or CGRP drive CeC activation during opioid withdrawal, and contribute to withdrawal-induced aversion as measured by place avoidance. In contrast to CeC, IPACc neurons lack direct PB inputs, and their activation is only modestly influenced by the PB, likely through indirect pathways. Furthermore, simultaneous inhibition of PB glutamatergic neurons and posterior striatal area neurons that include IPACc neurons robustly suppressed withdrawal-induced avoidance more strongly than inhibitions of each pathway alone, suggesting cooperative contribution of two largely separated neural systems in driving opioid withdrawal aversion.

### Neural pathway of CeC and IPACc activation during opioid withdrawal

Opioid withdrawal induces activation of neurons across many brain areas, with CeC and IPACc being two of the most strongly activated regions in the forebrain [5,6]. How these activations are triggered had not been well understood, but our data revealed that PB inactivation nearly silences withdrawal-induced CeC c-Fos suggesting that influences from other afferents, such as cortex, amygdala and thalamus [44–46], or local MORs [4] are not sufficient to compensate for the loss of PB input.

The PB contribution to withdrawal-induced CeC activation varies in a cell-type specific manner. Although CGRP^PB^ inhibition decreased c-Fos activation in CeC, consistent with its projection to CeC, the level of suppression became stronger with MOR^PB^ inhibition, and we obtained the most dramatic reductions withVG2^PB^ inhibition. This suggests that glutamatergic PB neurons other than CGRP^PB^ or MOR^PB^ also contribute to CeC activation, but further study is needed to identify these subtypes.

Unlike CeC, the IPACc region lacks direct input from PB neurons. The current study did not determine what brain structures contribute to IPACc activation during withdrawal, but IPACc receives inputs from insular cortex, midline thalamus and the midbrain [20,44,45,47,48]. In future studies, these afferents, as well as local MOR expression, can be examined as potential sources of excitatory drive for IPACc activation. Interestingly, despite the absence of direct projection from the PB, IPACc neuronal activation was partially attenuated by MOR^PB^ inhibition, possibly by multisynaptic interactions through intermediary regions or neuromodulatory mechanisms.

### Neural populations related to physical symptoms and aversiveness during opioid withdrawal

While acute opioid withdrawal in rodents produces somatic symptoms, such as jumping and wet dog shakes [49], it has remained unclear whether these signs are indicative of aversive states per se. Our study measured one widely studied physical sign, jumping, along with CPA, a readout of affective aspects of withdrawal. In our results, inhibition of CGRP^PB^ or MOR^PB^ during withdrawal reduced CPA without affecting jumping, while VG2^PB^ inhibition almost eliminated jumping, but only modestly reduced CPA, and finally IPACc area inhibition decreased CPA, with increased jumping. These dissociations between jumping and CPA suggest that jumping is not necessarily linked to the level of aversiveness.

In human, opioid withdrawal involves a diverse array of aversive somatic symptoms, including body aches, gastrointestinal upset and nausea [1]. In rodent studies, similar aversive conditions caused by noxious stimuli, such as footshock, LiCl and LPS, activate CeC-projecting CGRP^PB^ and/or CeC neurons [9,14,46]. These findings, along with our results, are broadly consistent with PB-CeC engagement during noxious events including opioid withdrawal. Additionally, previous research linking CeC to avoidance behaviors suggests that PB-mediated CeC activation could be associated with withdrawal aversion [9,46]. While our study did not directly inhibit CeC, our chemogenetic inhibition of PB neuron subsets, particularly VG2^PB^, produced strong suppression of CeC activation during withdrawal. However, VG2^PB^ inhibition produced only a modest reduction in withdrawal-induced CPA, suggesting that blocking of CeC activation is not sufficient to suppress the majority of withdrawal-induced aversion. Related to the role of CeA neurons in opioid withdrawal, a recent study demonstrated an aversive nature of CeA MOR+ neurons and their roles in physical symptoms of fentanyl withdrawal, but did not examine their involvement in withdrawal-induced aversion more directly, e.g. using CPA [50]. Our data on suppression of CeC activation via PB manipulations, suggests that CeC neurons are not the major contributor to withdrawal aversion. However, because we did not directly inhibit CeC or measure activity aside from c-Fos, one caveat is that residual CeC activity after PB inhibition could sustain withdrawal aversion.

While our data suggests that PB’s excitatory influence on CeC may not be the major contributor to withdrawal-induced aversion, it remains possible that other PB subcircuits play such roles. CGRP^PB^ neurons are involved in various pain and avoidance responses [9,14,15,51]. In our data, although CGRP^PB^ inhibition reduced withdrawal-induced CPA, it did so to a surprisingly modest degree. In contrast, inhibitions of MOR^PB^, which are more widespread in the PB [36,52], produced more significant reduction of CPA. Several neuron types in PBdl region, such as PDyn+ and Tacr1+ neurons, that produce aversion upon activation [18,37,38] and partially overlap with MOR^PB^ neurons [36,52], could be participating in withdrawal aversion. The modest reduction of CPA by VG2^PB^ inhibition suggests that VG2^PB^ may contain subpopulations of neurons that counteract the effects of other subsets on withdrawal aversion. That is, while MOR^PB^ contribute to withdrawal-induced aversiveness, some other subtypes of VG2^PB^ may drive opposite effects, as supported by prior reports of PB neuron subtypes activated by rewarding stimuli [39,40].

IPACc neurons are robustly activated during opioid withdrawal but have been largely ignored outside of a single study several decades ago [6]. Our data suggests that neurons in or near IPACc contribute to withdrawal aversion, particularly by augmenting withdrawal-related parabrachial activity. Despite its proximity to CeC, IPACc connectivity is more akin to the striatopallidal system, as it projects to GP, SNR [19], while receiving dense dopaminergic projections from midbrain [20]. Furthermore, IPACc neurons are not well connected to other EA structures, such as BNST and substantia innominate [19,21–23,46]. These anatomical characteristics suggest that IPACc operates within the basal ganglia system, which is distinct from the PB-CeC pathway. While it is unclear how IPACc-associated circuits would contribute to withdrawal aversion, its anatomical connection with insular cortex suggests the intriguing possibility of involvement in aversive visceral processes. However, it is also notable that our results do not rule out contribution of other nearby areas, such as the anterior tip of CEA and the posterior striatal region dorsal to IPACc, which were affected by hM4Di spread. Notably, while the striatal “tail’ has recently been implicated in avoidance behavior [53], that region is considerably posterior to the striatal regions involved here, suggesting a novel role for IPACc and adjacent regions that require further study to fully elucidate.

### Contribution of multiple circuits to distress of opioid withdrawal

Opioid withdrawal is a complex condition with a variety of aversive symptoms that may recruit multiple aversive circuits. Previous studies implicated several brain neuronal populations in driving the aversiveness of opioid withdrawal, including the PVT-accumbens shell pathway, medial habenula (MHb), and BNST [30,54,55]. Some of these regions, such as PVT and BNST, are anatomically connected with PB and IPACc, suggesting potential functional connections [13,45]. Further elucidation of the neuron populations that contribute to withdrawal aversion, as well as synaptic and functional connectivity between them would be helpful in developing strategies for reducing the aversiveness of opioid withdrawal via manipulations that target multiple appropriate systems, such as the PB-CeC and posterior striatal area in the current study.

## Acknowledgments

We would like to thank Calva Coleman, John Gibbons, Beyonce Getachew for technical help. We thank Yavin Shaham and Leandro Vendruscolo for scientific advice.

## Data Availability Statement

All data supporting the current study will be available upon request.

## Author contributions

Conceptualization: S.L. and S.I.

Investigation: S.L., K.S., J.R., J.C.

Data analysis: S.L., K.S., J.R.

Writing : S.L., T.J., S.I.

## Funding

This research was supported by the Intramural Research Program of the National Institute on Drug Abuse, National Institutes of Health (NIH). The contributions of the NIH authors are considered Works of the United States Government. The findings and conclusions presented in this paper are those of the author(s) and do not necessarily reflect the views of the NIH or the U.S. Department of Health and Human Services.

National Institutes of Health grant R37DA054370

Department of Neurobiology, University of Maryland Baltimore

## References

1. Kosten TR, Baxter LE. Review article: Effective management of opioid withdrawal symptoms: A gateway to opioid dependence treatment. American Journal on Addictions. 2019;28:55– 62.

2. Welsch L, Bailly J, Darcq E, Kieffer BL. The Negative Affect of Protracted Opioid Abstinence: Progress and Perspectives From Rodent Models. Biological Psychiatry. 2020;87:54–63.

3. Monroe SC, Radke AK. Opioid withdrawal: role in addiction and neural mechanisms. Psychopharmacology. 2023;240:1417–1433.

4. Mansour A, Fox CA, Burke S, Meng F, Thompson RC, Akil H, et al. Mu, delta, and kappa opioid receptor mRNA expression in the rat CNS: an in situ hybridization study. The Journal of Comparative Neurology. 1994;350:412–438.

5. Frenois F, Cador M, Caillé S, Stinus L, Le Moine C. Neural correlates of the motivational and somatic components of naloxone-precipitated morphine withdrawal. The European Journal of Neuroscience. 2002;16:1377–1389.

6. Veinante P, Stoeckel ME, Lasbennes F, Freund-Mercier MJ. c-Fos and peptide immunoreactivities in the central extended amygdala of morphine-dependent rats after naloxone-precipitated withdrawal. European Journal of Neuroscience. 2003;18:1295–1305.

7. Koob GF. A Role for Brain Stress Systems in Addiction. Neuron. 2008;59:11–34.

8. Alheid GF. Extended amygdala and basal forebrain. Annals of the New York Academy of Sciences, vol. 985, New York Academy of Sciences; 2003. p. 185–205.

9. Han S, Soleiman M, Soden M, Zweifel L, Palmiter RD. Elucidating an Affective Pain Circuit that Creates a Threat Memory. Cell. 2015;162:363–374.

10. Neugebauer V, Li W, Bird GC, Han JS. The amygdala and persistent pain. Neuroscientist. 2004;10:221–234.

11. Bernard JF, Besson JM. The spino(trigemino)pontoamygdaloid pathway: Electrophysiological evidence for an involvement in pain processes. Journal of Neurophysiology. 1990;63:473– 490.

12. Huang D, Grady FS, Peltekian L, Laing JJ, Geerling JC. Efferent projections of CGRP/Calca-expressing parabrachial neurons in mice. Journal of Comparative Neurology. 2021;529:2911– 2957.

13. Huang D, Grady FS, Peltekian L, Geerling JC. Efferent projections of Vglut2, Foxp2, and Pdyn parabrachial neurons in mice. The Journal of Comparative Neurology. 2021;529:657–693.

14. Campos CA, Bowen AJ, Roman CW, Palmiter RD. Encoding of danger by parabrachial CGRP neurons. Nature. 2018;555:617–620.

15. Bowen AJ, Chen JY, Huang YW, Baertsch NA, Park S, Palmiter RD. Dissociable control of unconditioned responses and associative fear learning by parabrachial cgrp neurons. eLife. 2020;9:1–50.

16. Barik A, Thompson JH, Seltzer M, Ghitani N, Chesler AT. A Brainstem-Spinal Circuit Controlling Nocifensive Behavior. Neuron. 2018;100:1491–1503.e3.

17. Chiang MC, Nguyen EK, Canto-Bustos M, Papale AE, Oswald A-MM, Ross SE. Divergent Neural Pathways Emanating from the Lateral Parabrachial Nucleus Mediate Distinct Components of the Pain Response. Neuron;0.

18. Deng J, Zhou H, Lin JK, Shen ZX, Chen WZ, Wang LH, et al. The Parabrachial Nucleus Directly Channels Spinal Nociceptive Signals to the Intralaminar Thalamic Nuclei, but Not the Amygdala. Neuron. 2020;107:909–923.e6.

19. Shammah-Lagnado SJ, Alheid GF, Heimer L. Striatal and central extended amygdala parts of the interstitial nucleus of the posterior limb of the anterior commissure: evidence from tract-tracing techniques in the rat. The Journal of Comparative Neurology. 2001;439:104–126.

20. Yamaguchi T, Ehara A, Nakadate K, Ueda S. Tyrosine hydroxylase afferents to the interstitial nucleus of the posterior limb of the anterior commissure are neurochemically distinct from those projecting to neighboring nuclei. Journal of Chemical Neuroanatomy. 2018;90:98–107.

21. Dong HW, Swanson LW. Projections from the rhomboid nucleus of the bed nuclei of the stria terminalis: implications for cerebral hemisphere regulation of ingestive behaviors. The Journal of Comparative Neurology. 2003;463:434–472.

22. Bourgeais L, Gauriau C, Bernard JF. Projections from the nociceptive area of the central nucleus of the amygdala to the forebrain: a PHA-L study in the rat. The European Journal of Neuroscience. 2001;14:229–255.

23. Shammah-Lagnado SJ, Alheid GF, Heimer L. Afferent connections of the interstitial nucleus of the posterior limb of the anterior commissure and adjacent amygdalostriatal transition area in the rat. Neuroscience. 1999;94:1097–1123.

24. Liu HM, Liao ML, Liu GX, Wang LJ, Lian D, Ren J, et al. IPAC integrates rewarding and environmental memory during the acquisition of morphine CPP. Science Advances. 2023;9:eadg5849.

25. Furlan A, Corona A, Boyle S, Sharma R, Rubino R, Habel J, et al. Neurotensin neurons in the extended amygdala control dietary choice and energy homeostasis. Nature Neuroscience. 2022;25:1470–1480.

26. Chang S, Fermani F, Lao CL, Huang L, Jakovcevski M, Di Giaimo R, et al. Tripartite extended amygdala-basal ganglia CRH circuit drives locomotor activation and avoidance behavior. Science Advances. 2022;8:eabo1023.

27. Tanaka DH, Li S, Mukae S, Tanabe T. Genetic Access to Gustatory Disgust-Associated Neurons in the Interstitial Nucleus of the Posterior Limb of the Anterior Commissure in Male Mice. Neuroscience. 2019;413:45–63.

28. Kesner AJ, Shin R, Calva CB, Don RF, Junn S, Potter CT, et al. Supramammillary neurons projecting to the septum regulate dopamine and motivation for environmental interaction in mice. Nat Commun. 2021;12:2811.

29. Warren BL, Kane L, Venniro M, Selvam P, Quintana-Feliciano R, Mendoza MP, et al. Separate vmPFC Ensembles Control Cocaine Self-Administration Versus Extinction in Rats. J Neurosci. 2019;39:7394–7407.

30. Boulos LJ, Ben Hamida S, Bailly J, Maitra M, Ehrlich AT, Gavériaux-Ruff C, et al. Mu opioid receptors in the medial habenula contribute to naloxone aversion. Neuropsychopharmacology : Official Publication of the American College of Neuropsychopharmacology. 2020;45:247–255.

31. Chamberlin NL, Mansour A, Watson SJ, Saper CB. Localization of mu-opioid receptors on amygdaloid projection neurons in the parabrachial nucleus of the rat. Brain Research. 1999;827:198–204.

32. Palmiter RD. The Parabrachial Nucleus: CGRP Neurons Function as a General Alarm. Trends in Neurosciences. 2018;41:280–293.

33. Kovács KJ. c-Fos as a transcription factor: A stressful (re)view from a functional map. Neurochemistry International. 1998;33:287–297.

34. Weinberg MS, Girotti M, Spencer RL. Restraint-induced fra-2 and c-fos expression in the rat forebrain: Relationship to stress duration. Neuroscience. 2007;150:478–486.

35. Cullity ER, Guerin AA, Perry CJ, Kim JH. Examining Sex Differences in Conditioned Place Preference or Aversion to Methamphetamine in Adolescent and Adult Mice. Front Pharmacol. 2021;12:770614.

36. Pauli JL, Chen JY, Basiri ML, Park S, Carter ME, Sanz E, et al. Molecular and anatomical characterization of parabrachial neurons and their axonal projections. eLife. 2022;11:e81868.

37. Luskin AT, Bhatti DL, Mulvey B, Pedersen CE, Girven KS, Oden-Brunson H, et al. Extended amygdala-parabrachial circuits alter threat assessment and regulate feeding. Science Advances. 2021;7:eabd3666.

38. Kim DY, Heo G, Kim M, Kim H, Jin JA, Kim HK, et al. A neural circuit mechanism for mechanosensory feedback control of ingestion. Nature. 2020;580:376–380.

39. Fu O, Iwai Y, Kondoh K, Misaka T, Minokoshi Y, Nakajima K ichiro. SatB2-Expressing Neurons in the Parabrachial Nucleus Encode Sweet Taste. Cell Reports. 2019;27:1650–1656.e4.

40. Jarvie BC, Chen JY, King HO, Palmiter RD. Satb2 neurons in the parabrachial nucleus mediate taste perception. Nature Communications. 2021;12:224.

41. Lisman J, Schulman H, Cline H. The molecular basis of CaMKII function in synaptic and behavioural memory. Nature Reviews Neuroscience. 2002;3:175–190.

42. Fukunaga K, Goto S, Miyamoto E. Immunohistochemical Localization of Ca2+/Calmodulin-Dependent Protein Kinase II in Rat Brain and Various Tissues. Journal of Neurochemistry. 1988;51:1070–1078.

43. Allen Institute for Brain Science. Allen Mouse Brain Atlas ISH. 2004.

44. McDonald AJ, Shammah-Lagnado SJ, Shi C, Davis M. Cortical afferents to the extended amygdala. Annals of the New York Academy of Sciences. 1999;877:309–338.

45. Li S, Kirouac GJ. Projections from the paraventricular nucleus of the thalamus to the forebrain, with special emphasis on the extended amygdala. The Journal of Comparative Neurology. 2008;506:263–287.

46. Bowen AJ, Huang YW, Chen JY, Pauli JL, Campos CA, Palmiter RD. Topographic representation of current and future threats in the mouse nociceptive amygdala. Nature Communications. 2023;14:196.

47. Otake K, Nakamura Y. Forebrain neurons with collateral projections to both the interstitial nucleus of the posterior limb of the anterior commissure and the nucleus of the solitary tract in the rat. Neuroscience. 2003;119:623–628.

48. Vertes RP, Hoover WB, Rodriguez JJ. Projections of the central medial nucleus of the thalamus in the rat: Node in cortical, striatal and limbic forebrain circuitry. Neuroscience. 2012;219:120–136.

49. Alvarez-Bagnarol Y, Vendruscolo LF, Marchette RCN, Francis C, Morales M. Neuronal Correlates of Hyperalgesia and Somatic Signs of Heroin Withdrawal in Male and Female Mice. eNeuro. 2022;9:ENEURO.0106-22.2022.

50. Chaudun F, Python L, Liu Y, Hiver A, Cand J, Kieffer BL, et al. Distinct µ-opioid ensembles trigger positive and negative fentanyl reinforcement. Nature. 2024;630:141–148.

51. Kang SJ, Liu S, Ye M, Kim DI, Pao GM, Copits BA, et al. A central alarm system that gates multi-sensory innate threat cues to the amygdala. Cell Reports. 2022;40:111222.

52. Liu S, Ye M, Pao GM, Song SM, Jhang J, Jiang H, et al. Divergent brainstem opioidergic pathways that coordinate breathing with pain and emotions. Neuron. 2022;110:857–873.e9.

53. Tsutsui-Kimura I, Tian ZM, Amo R, Zhuo Y, Li Y, Campbell MG, et al. Dopamine in the tail of the striatum facilitates avoidance in threat-reward conflicts. Nat Neurosci. 2025;28:795– 810.

54. Zhu Y, Wienecke CFR, Nachtrab G, Chen X. A thalamic input to the nucleus accumbens mediates opiate dependence. Nature. 2016;530:219–222.

55. Delfs JM, Zhu Y, Druhan JP, Aston-Jones G. Noradrenaline in the ventral forebrain is critical for opiate withdrawal-induced aversion. Nature. 2000;403:430–434.

56. Paxinos G, Franklin KBJ. The Mouse Brain in Sterotaxic Coordinates. Elsevier/Academic Press; 2013.

