## Supplemental Information for "Parabrachial-amygdala circuit cooperates with a posterior striatal area to drive opioid withdrawal aversion"

Seung-Chan Lee *et al.*

**This PDF file includes:**

Figures S1 to S6
Table S1
Supplementary Materials and Methods

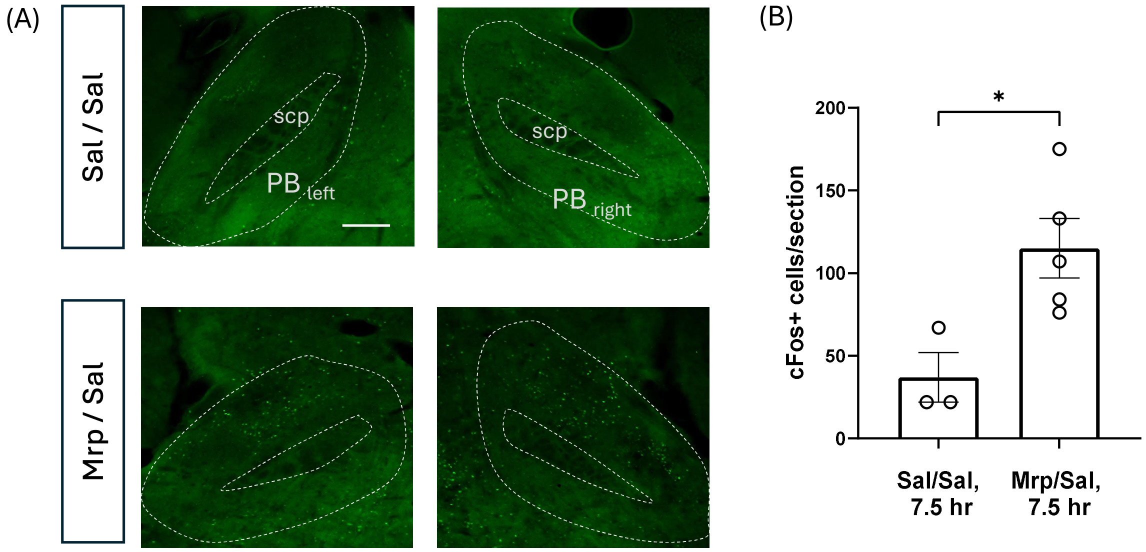


**Figure S1. Increase of c-Fos expression in the PB after morphine treatment**
**(A)** Coronal sections showing representative c-Fos expressions of the mice that received either saline or morphine twice a day for 6 consecutive days. On day 6, 6 hours after the morning saline/morphine treatment, a single saline injection was given. 1.5 hours after this saline injection, brains were extracted for IHC. Scale bar, 200 µm. **(B)** Numbers of c-Fos+ neurons per section in the PB of the Sal/Sal and Mrp/Sal groups. * $p < 0.05$ (two-tailed t-tests)

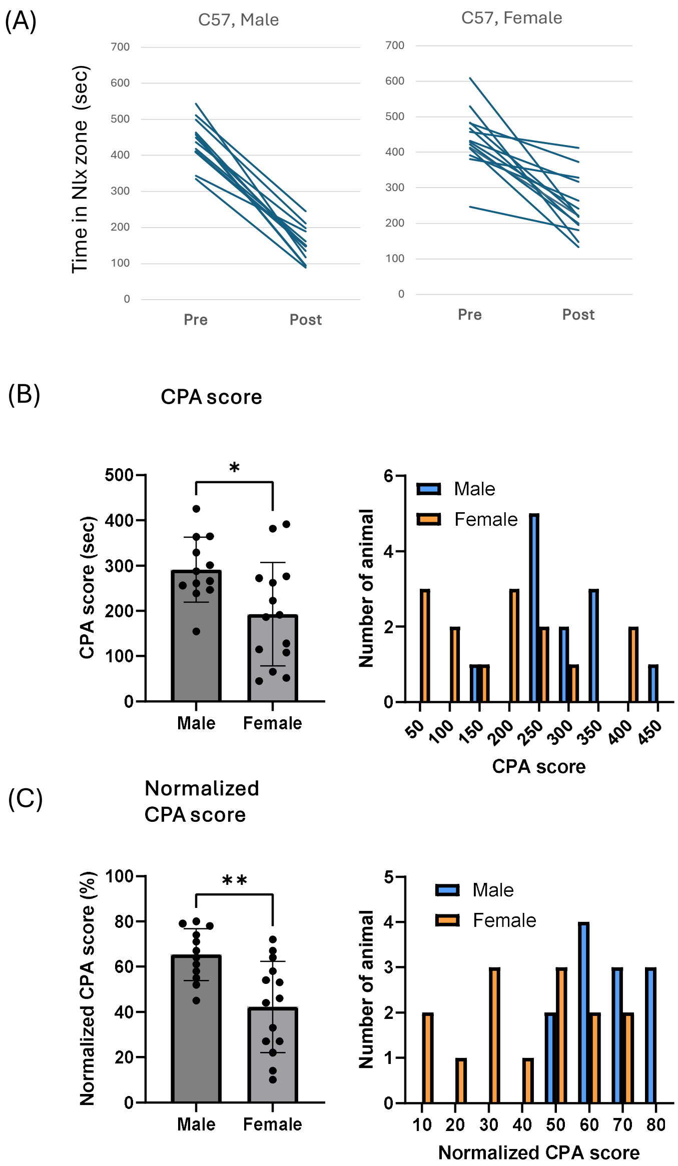


**Figure S2. Sex differences in opioid withdrawal-induced conditioned place avoidance**
Male and female C57 WT mice were treated with morphine followed by naloxone precipitated withdrawal as described in Figure 2A. **(A)** Time spent in naloxone-paired zone before and after the conditioning of male (left) and female (right) mice. **(B-C)** Precipitated withdrawal produced stronger CPA in male mice than female mice as indicated by CPA scores (B) and normalized CPA scores (C). * $p < 0.05$, ** $p < 0.01$ (two-tailed t-tests)

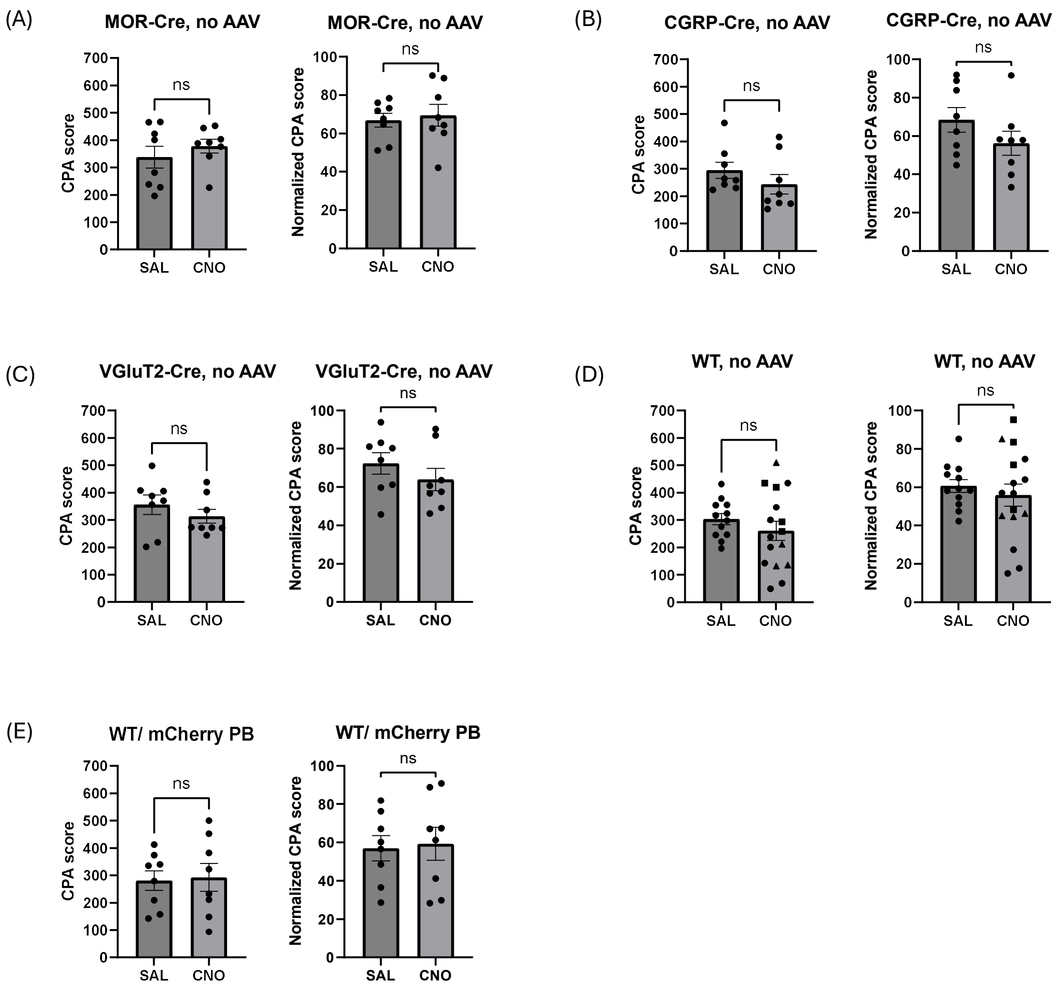


**Figure S3. CNO treatments in the absence of hM4Di expression do not affect withdrawal-induced CPA.**

The effect of CNO treatment on withdrawal-induced CPA was measured in mice that do not express hM4Di. Mice were treated with morphine followed by naloxone as described in Figure 2A. We examined effect of CNO in 4 mouse lines that received no surgery (A-D). To shed light on possible dose effect of CNO, the WT naïve mice received varied CNO doses: 4 mg/kg, triangles; 5 mg/kg, circles; 6 mg/kg, squares (D). Some WT mice received bilateral intra-PB injections of AAV-hsyn-mCherry (E) and were treated with 5 mg/kg CNO. ns: $p > 0.05$ by two-sided t-test.

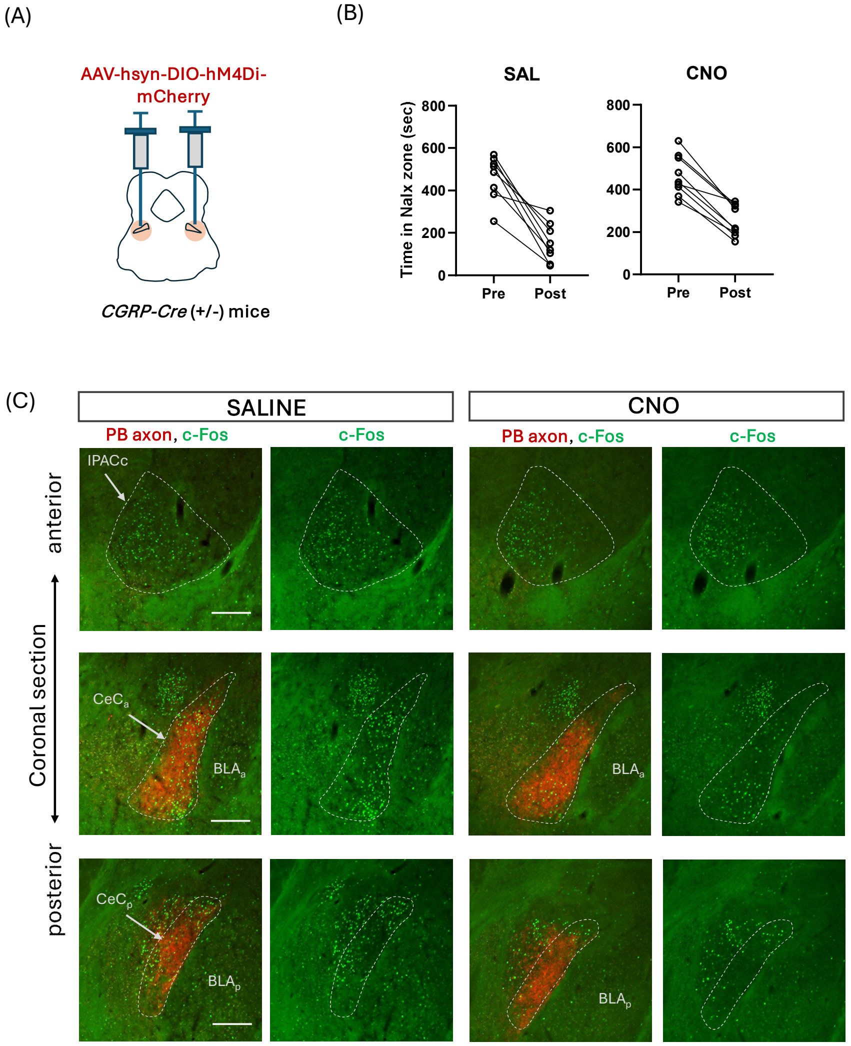


**Figure S4. Chemogenetic inhibition of CGRP$^{PB}$ affects opioid withdrawal-induced place avoidance and c-Fos counts in CeC.**

(A) CGRP-Cre mice received bilateral injection of an AAV-hsyn-DIO-hM4Di-mCherry into the PB. (B) Time spent in naloxone-paired compartment before and after conditioning. See Fig. 3D for CPA scores. (C) Coronal sections showing representative c-Fos expression of saline- and CNO-treated mice. See Fig. 3E for group data. Abbreviations: $CeC_a$, anterior capsular central amygdala; $CeC_p$, posterior capsular central amygdala; $BLA_a$, anterior basolateral amygdala; $BLA_p$: posterior basolateral amygdala. Scale bar 200 μm.

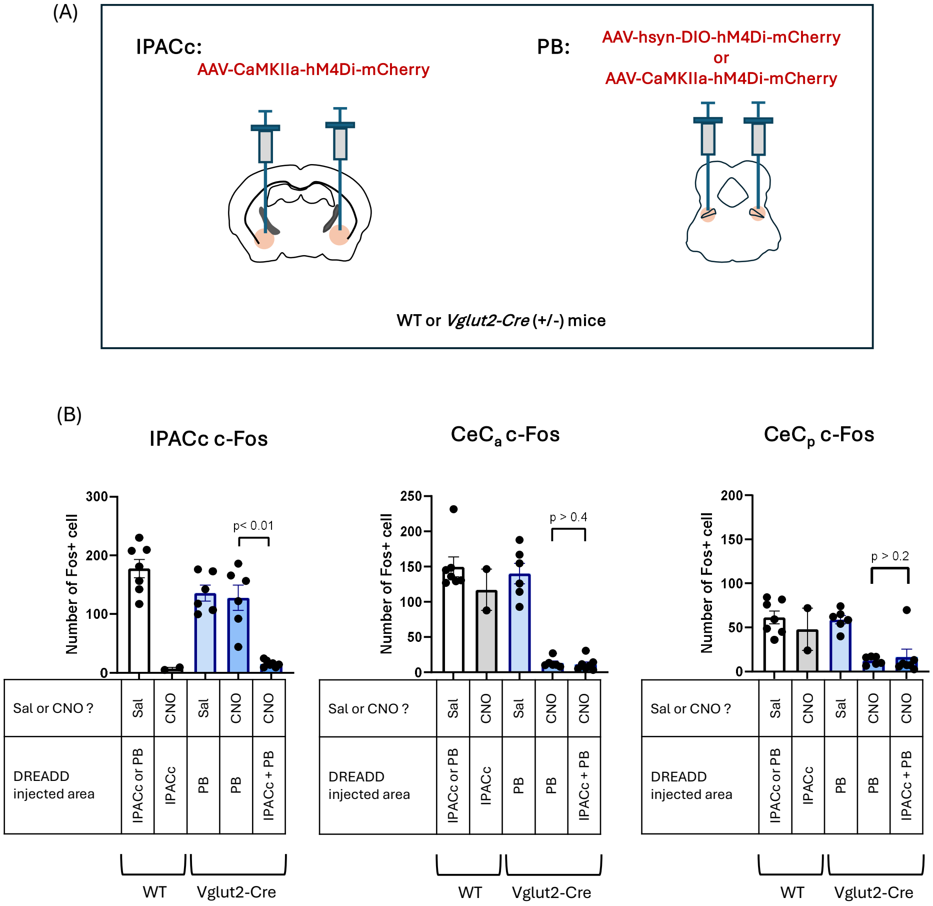


**Figure S5. Chemogenetic inhibition of IPACc area neurons using AAV-CaMKIIα-hM4Di suppresses withdrawal-induced c-Fos expression in IPACc neurons.**

**(A)** WT mice received AAV-CaMKIIα-hM4Di into the IPACc area (3 mice), or AAV-CaMKIIα-hM4Di into the PB (6 mice). Vglut2-Cre mice received AAV-CaMKIIα-hM4Di into the IPACc area and AAV-DIO-hM4Di into the PB (dual injection, 8 mice), or AAV-DIO-hM4Di into the PB with no injections into the IPACc (12 mice). **(B)** c-Fos expressions were examined after precipitated withdrawal in the IPACc (left graph), $CeC_a$ (middle graph), and $CeC_p$ (right graph). Prior to withdrawal induction, three groups received CNO, while two groups received saline as described in the table. IPACc c-Fos expression was strongly suppressed in mice with IPACc hM4Di and CNO treatment. $CeC_a$, anterior CeC; $CeC_p$, posterior CeC.

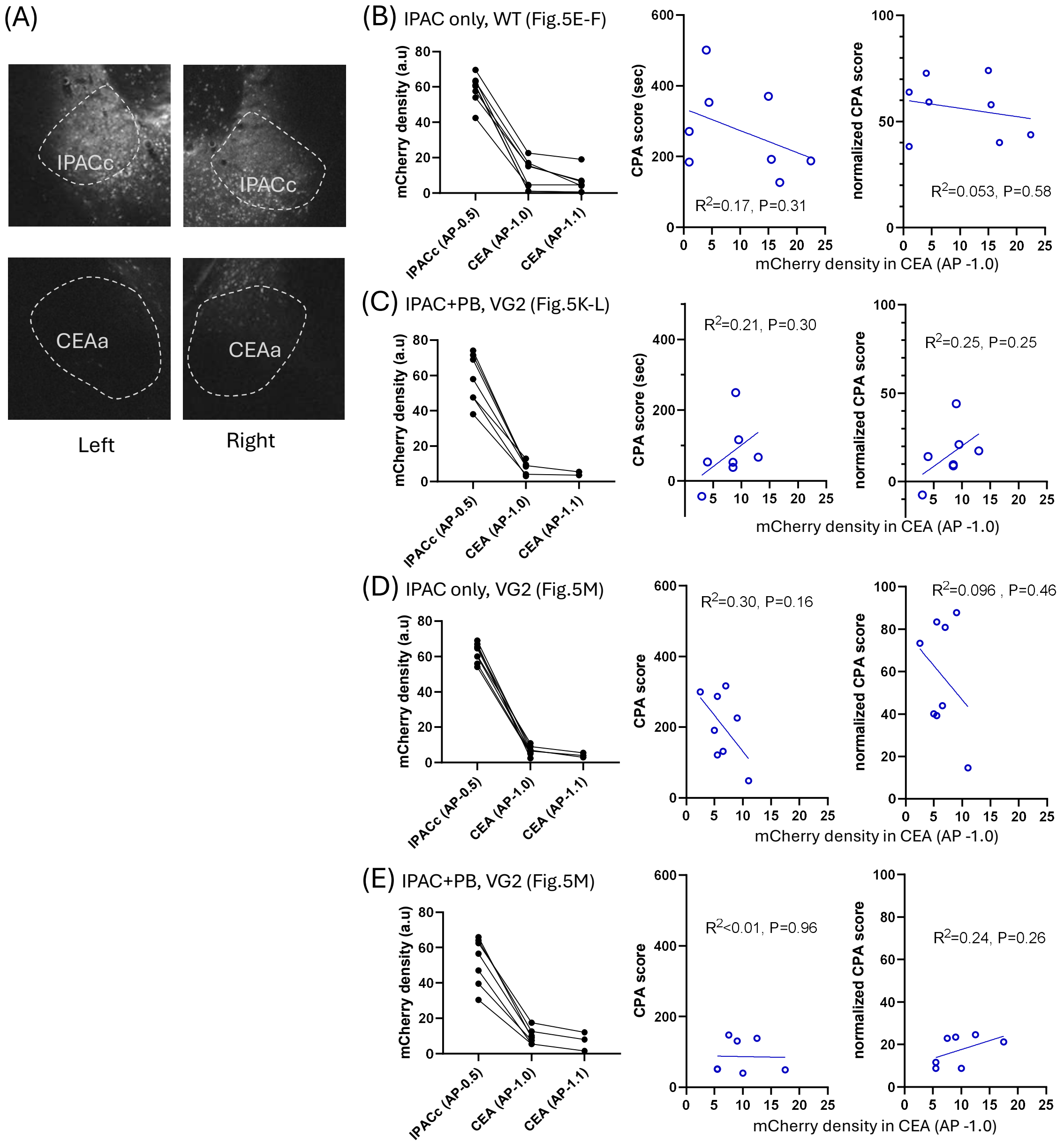


**Figure S6. Limited spread of hM4Di expression into CEA after IPACc area AAV injection.**

**(A)** An example of hM4Di-mCherry expression in IPACc area and anterior portion of CEA (AP -1.0 position of Paxinos mouse brain atlas) after injection of AAV-CaMKIIα-hM4Di-mCherry into the IPACc area of WT mice.

**(B)** Left: Optical density of hM4Di-mCherry signal in IPACc (AP -0.5), and two AP levels of CEA (AP -1.0, AP -1.1), of WT mice that received AAV-CaMKIIα-hM4Di-mCherry injection into IPACc area (the same mice used in Fig. 5E-F). AP -1.0 corresponds to ~300 um posterior position from the anterior tip of CEA. mCherry density is mean value of the measurements from left and right hemispheres. Some animals have only IPACc and CEA (AP -1.0) measurements. Middle, Right: Linear correlation analyses between mCherry signal in CEA (AP -1.0) and CPA score (Middle) or normalized CPA score (Right).

**(C)** The same analyses as (B) were done in Vglut2-cre mice that received AAV-CaMKIIα-hM4Di-mCherry injection into IPACc area and AAV-syn-DIO-hM4Di-mCherry injection into PB (the same mice used in Fig. 5K-L).
**(D)** The same analyses as (B) were done in Vglut2-cre mice that received AAV-CaMKIIα-hM4Di-mCherry injection into IPACc area (the same mice used in Fig. 5M).
**(E)** The same analyses as (B) were done in Vglut2-cre mice that received AAV-CaMKIIα-hM4Di-mCherry injection into IPACc area and AAV-syn-DIO-hM4Di-mCherry injection into PB (the same mice used in Fig. 5M).

### Supplementary Materials and Methods

#### Reagents

| Reagent or Resource | Source | Identifier |
|---|---|---|
| AAV9-CamKIIα-hM4Di-mCherry | Addgene | 50477-AAV9 |
| AAV9-hsyn-DIO-hM4Di-mCherry | Addgene | 44362-AAV9 |
| AAV9-CamKIIα-mCherry | Addgene | 114469-AAV9 |
| RNAscope Probe: *Oprm1* | Advance Cell Diagnostics | 315841 |
| RNAscope Probe: *Calca* | Advance Cell Diagnostics | 578771 |
| RNAscope Probe: *cfos* | Advance Cell Diagnostics | 316921 |
| RNAscope Probe: *Camk2a* | Advance Cell Diagnostics | 445231 |
| RNAscope Probe: *Slc17a6 (vGlut2)* | Advance Cell Diagnostics | 319171 |

#### Animals

C57BL/6J mice (Strain #: 000664) were purchased from Jackson Laboratory. Vglut2-Cre (strain #: 016963) and CGRP-Cre (strain #: 033168) homozygote mice were purchased from Jackson Laboratory and crossed with C57BL/6J mice to generate Vglut2-Cre and CGRP-Cre heterozygote mice. MOR-Cre mice were provided by B. L. Kieffer and crossed with C57BL/6J mice to generate MOR-Cre heterozygote mice. Mice were 4-10 months old at the time of the first withdrawal induction. We observed sex differences in opioid withdrawal-induced conditioned place avoidance in a pilot experiment (Supplementary Fig. 2). Therefore, only male mice were used for this study. Mice were group-housed and maintained on a normal 12-hour light/dark cycle (lights on at 7 am) with food and water ad libitum. All protocols for animal experiments were approved by the Animal Care and Use Committee of the National Institute on Drug Abuse.

#### Drugs

Morphine sulfate pentahydrate ($(C_{17}H_{19}NO_3)_2 \cdot H_2SO_4 \cdot 5H_2O$, molecular weight 758.8) was obtained as an aqueous solution from NIDA pharmacy. Naloxone hydrochloride dihydrate was purchased from Sigma Aldrich (N7758, $C_{19}H_{21}NO_4 \cdot HCl \cdot 2H_2O$, molecular weight 399.9). Clozapine-N-oxide dihydrochloride (CNO; $C_{19}H_{19}ClN_4O \cdot 2HCl$, molecular weight 415.74) was purchased from Hello Bio (HB6149). CNO was dissolved in water and stored at -20°C for up to 30 days.

**Stereotaxic surgery for virus injection**

Under isoflurane anesthesia (maintenance at 1.5-2%), mice were placed on a stereotaxic frame (David Kopf Instruments). The skull was exposed, and holes were produced with a micromotor drill. A pulled glass pipette was backfilled with virus solution and lowered to the target area. Viruses were pressure-injected (90 nl for the PBN and 50 nl for the IPACc) at a rate of ~0.05 µL/min, except that AAV-CamKIIα-mCherry injection into the PBN was 180 nl. Coordinates for virus injections were: PBN (AP, -5.25 mm; ML, ±1.5 mm; DV, -3.8 mm); IPACc (AP, -0.45 mm; ML, ±2.75 mm; DV, -4.7 mm). After surgery, mice were single-housed for 2 weeks to protect sutures and then group-housed with previous cage mates that received a similar surgery.

AAVs were diluted with sterile saline (0.9% sodium chloride) on the day of use. The titers (genome copy/ml) of adeno-associated viruses (AAVs) were 5.0 x $10^{12}$ for AAV9-CamKIIα-hM4Di-mCherry, 5.0 x $10^{12}$ for AAV9-CamKIIα-mCherry, and 1.0 x $10^{13}$ for AAV9-hsyn-DIO-hM4Di-mCherry.

**Naloxone-precipitated morphine withdrawal**

To induce morphine dependence, mice were injected subcutaneously twice a day with escalating doses of morphine sulfate pentahydrate (20, 20, 40, 40, 60, 60, 80, 80, 100, 100 mg/kg body weight) over six consecutive days (Figure 1A). Six hours after the last morning morphine injection on the 6th day, half of the mice subcutaneously received naloxone hydrochloride dihydrate (0.19 mg/kg), while the other half received saline (4 ml/kg).

**Detection of c-Fos expression**

Ninety minutes after saline or naloxone injection, mice were deeply anesthetized with isoflurane and transcardially perfused with 10 ml of phosphate-buffered saline (PBS) followed by 20 ml of paraformaldehyde (PFA, 4%). Brains were extracted and postfixed in 4% PFA at 4°C for 24 hours, then transferred to 30% sucrose for 48 hours for cryoprotection at 4°C, and frozen with dry ice for storage at -80°C. The frozen brains were sectioned at 50 µm on a cryostat (Leica Microsystems). The sections were rinsed in PBS 3 times for 5 minutes and incubated for 1 hour in blocking solution containing 0.30% Triton X-100 and 5% normal donkey serum. Then, they were incubated for 20 hours with primary antibody (1:800, Cell Signaling Technology, Phospho-c-Fos (Ser32) (D82C12) XP® Rabbit mAb #5348), rinsed in PBS 4 times for 5 minutes each, followed by a 2-hour incubation with secondary antibody (1:800, Jackson ImmunoResearch Laboratories, Alexa Fluor® 488 AffiniPure™ $F(ab')_2$ Fragment Donkey Anti-Rabbit IgG (H+L)). After being rinsed in PBS 3 times for 5 minutes, the sections were mounted on micro slides superfrost plus (VWR, 48311-703) and coverslipped with Fluoromount-G mounting medium (ThermoFisher). Images were captured on a Keyence BZ-X at 10x resolution. The stained cells were semi-automatically quantified using QuPath software [53].

**RNAscope in situ hybridization**

Thirty-five minutes after saline or morphine injection, animals were anesthetized, and brains were quickly removed and frozen in isopentane chilled with dry ice and stored in a tightly sealed container at -80°C. The brains were sectioned in 16 µm thickness, and sections were mounted onto Superfrost plus slides (VWR, 48311-703), then dried for 60 minutes at -20°C and stored in tightly sealed bags at -80°C until use.

For RNAscope in situ hybridization, RNAscope fluorescence multiplex v1 and v2 kits (Advanced Cell Diagnostics) were used. The sections on slides were incubated in 4% paraformaldehyde in PBS at 4°C for one hour and dehydrated in increasing concentrations (50%, 70%, and 100%) of ethanol for 5 minutes for each concentration. After drying at room temperature, the sections were pretreated with hydrogen peroxide for 10 minutes at room temperature. After washing twice with distilled water, the sections were incubated with protease for 30 minutes at room temperature, washed twice with distilled water, and hybridized with probes listed below for 2 hours at 40°C. For RNAscope multiplex fluorescent reagent kit v2, we used opal 520, opal 570, and opal 690 reagent packs. Images were captured on a confocal microscope (ZEISS LSM 880) using ZenBlue at 20x resolution. Cell counts and colocalization were determined semi-automatically using QuPath [53]. Probes used for staining are *Oprm1* (ACDBio 315841), *Calca* (ACDBio 578771), *c-fos* (ACDBio 316921), *CamK2a* (ACDBio 445231), *Slc17a6* (ACDBio 319171). Cells were classified as positive if they expressed three or more puncta for cell-type marker genes (CaMKIIα, MOR, CGRP, VGluT2). To identify *c-fos*+ activated neurons, cells that expressed ten or more *c-fos*+ fluorescent puncta were classified as positive.

**Procedure for the effects of inhibitory DREADD manipulations on locomotor activities, CPA, and c-Fos expression**

**Locomotor activity assay**: Four to 5 weeks after AAV injection surgery, locomotor activity assay was performed over three consecutive days (Fig. 2A). Locomotor activity was measured in mouse chambers (15.9 x 14.0 x12.7 cm, Med Associates, St. Albans, VT) equipped with 4 pairs of infrared detectors. On day 1, each mouse was placed in the chamber for 5 min for habituation. On day 2, mice received a saline injection (i.p) and 35 min later, were placed in the chambers for 15 min. On day 3, mice received an i.p. injection of CNO (5 mg/kg) and 35 min later, were placed in the chambers for 15 min. The number of IR beam breaks during 15 min was counted as readout of locomotor activity.

**Withdrawal-induced conditioned place avoidance**

CPA experiment was performed using a black plexiglass box apparatus (ANY-maze mouse place preference box, Stoelting Co, Wood Dale, IL) with left and right compartments (W18 x L20 x H35 cm) and a middle compartment (20 x 10 x 35 cm). The left and right compartments are distinguished by visual, tactile and olfactory cues: the left compartment had yellow vertical stripes on the wall and a white floor mat textured with grid pattern and was scented with a lemon extract. The right compartment had white circles on the wall and a perforated metal floor mat and was scented with a peppermint extract. As illustrated in Figure 2A, mice were habituated to the CPA box for about 5 min on day -1. On day 0, each mouse was placed in the middle compartment of the CPA box and was allowed to explore the entire box freely for 15 min. Time that mice spent in each compartment of the box was measured as pre-conditioning place values. Mice that displayed 230-750 sec range for the time spent in naloxone-paired side of the CPA box during baseline measurement were included for the CPA test. From the afternoon of day 1, mice received the morphine injections as described above in the Naloxone-precipitated morphine withdrawal section (Fig. 2A). On day 5, six hours after the morning morphine injection, mice subcutaneously received saline injection (3.3 ml/kg) and was confined in the left chamber of the CPA box for 20 min. On day 6, 5.5 hr after the morning morphine injection, mice received an injection of either CNO (5 mg/kg) or saline (4 ml/kg). Thirty minutes after the injection (6 hours after the morning morphine injection), mice received naloxone hydrochloride dihydrate (0.19 mg/kg) and was confined in the right side of the CPA box for 20 min for naloxone conditioning (also termed as 'withdrawal conditioning'). If naloxone component only (molecular weight 327.4) is considered (excluding hydrochloride counter anions and water molecules), naloxone dose was 0.156 mg per kg body weight. Withdrawal behavior was video recorded from the camera attached above the chamber. Jumping incidents during early 15 min were counted manually offline. On day 7, mice were placed in the middle compartment of the CPA box and allowed to explore the entire box freely for 15 min. The level of CPA was accessed by two ways: CPA score and normalized CPA score. CPA score was calculated by subtracting the time spent in the naloxone-paired compartment during the test (post-conditioning) from the time spent in the same side during baseline (pre-conditioning). Normalized CPA score was calculated by dividing CPA score with the time spent in the naloxone-paired compartment during baseline.

**c-Fos expression**: We sought to determine how the CNO treatment affected the c-Fos expression in the CeC and IPACc and used some of the mice that had gone through the CPA procedure described above. Starting at a few hours after the CPA test on day 7 (Figure 2A), they received additional morphine sulfate pentahydrate injections of 60, 80, 100, 100 mg/kg in this order over 3 consecutive days. On day 9, the mice received the injection procedure as described above in the Naloxone-precipitated morphine withdrawal section and then c-Fos procedure as described in the Detection of c-Fos section. The mice that received naloxone on day 8 again received naloxone on day 11.

**Statistical Analysis**

Statistical analyses were performed with GraphPad Prism. Statistical significance was taken as $*p < 0.05$, $**p < 0.01$, and $***p < 0.001$. Default statistical test is the Student's t test (paired and unpaired) and summary data are expressed as means ± SEM, when data passed normality test (Kolmogorov-Smirnov test, $p > 0.05$). In case that data did not pass normality test, non-parametric tests (Mann-Whiteney for unpaired test, Wilcoxon signed- rank test for paired test) were used and summary data are expressed as median ± IQR. All tests were two sided.
